# THINGSvision: a Python toolbox for streamlining the extraction of activations from deep neural networks

**DOI:** 10.1101/2021.03.11.434979

**Authors:** Lukas Muttenthaler, Martin N. Hebart

**Affiliations:** Vision and Computational Cognition Group, Max Planck Institute for Human Cognitive and Brain Sciences, Leipzig, Germany; Machine Learning Group, Technical University of Berlin, Germany

**Author notes:** Corresponding authors: Lukas, L.M., Muttenthaler;, Martin, M.N.H, Hebart. **Author roles:** Conceptualization: L.M. & M.N.H; Funding acquisition: M.N.H.; Resources: L.M. & M.N.H.; Software: L.M.; Supervision: M.N.H.; Visualization: L.M.; Writing – original draft: L.M.; Writing – final manuscript: L.M. & M.N.H.

**Keywords:** Deep Neural Networks, Computational Neuroscience, Computer Vision, Feature extraction, Artificial Intelligence, Python

## Abstract

Over the past decade, deep neural network (DNN) models have received a lot of attention due to their near-human object classification performance and their excellent prediction of signals recorded from biological visual systems. To better understand the function of these networks and relate them to hypotheses about brain activity and behavior, researchers need to extract the activations to images across different DNN layers. The abundance of different DNN variants, however, can often be unwieldy, and the task of extracting DNN activations from different layers may be non-trivial and error-prone for someone without a strong computational background. Thus, researchers in the fields of cognitive science and computational neuroscience would benefit from a library or package that supports a user in the extraction task. `THINGSvision` is a new Python module that aims at closing this gap by providing a simple and unified tool for extracting layer activations for a wide range of pretrained and randomly-initialized neural network architectures, even for users with little to no programming experience. We demonstrate the general utility of `THINGsvision` by relating extracted DNN activations to a number of functional MRI and behavioral datasets using representational similarity analysis, which can be performed as an integral part of the toolbox. Together, `THINGSvision` enables researchers across diverse fields to extract features in a streamlined manner for their custom image dataset, thereby improving the ease of relating DNNs, brain activity, and behavior, and improving the reproducibility of findings in these research fields.

## 1 Introduction

In recent years, deep neural networks (DNNs) have sparked a lot of interest in the connected fields of cognitive science, computational neuroscience, and artificial intelligence. This is mainly owing to their power as arbitrary function approximators (LeCun, Bengio, & Hinton, 2015), their near-human performance on object recognition and natural language understanding tasks (e.g., Russakovsky et al. (2015); Wang et al. (2019, 2018)), and, most crucially, the fact that their latent representations often show a close correspondence to brain recordings and behavioral measurements (Güçlü & van Gerven, 2014; Khaligh-Razavi & Kriegeskorte, 2014; Kietzmann, McClure, & Kriegeskorte, 2018; King, Groen, Steel, Kravitz, & Baker, 2019; Kriegeskorte, 2015; Schrimpf et al., 2018; Schrimpf, Kubilius, et al., 2020; Yamins et al., 2014).

One important limiting factor for a much broader interdisciplinary adoption of DNNs as computational models lies in the difficulty of extracting layer activations for DNNs. This difficulty is twofold. First, the number of existing models is enormous and increases by the day. Due to this diversity, an extraction strategy that is suited for one model may not apply to any other model. Second, for users without a strong programming background it can be non-trivial to extract features while being confident that no mistakes were made in the process, for example during image preprocessing, layer selection, or making sure that images corresponded to extracted activations. Beyond these difficulties, even experienced programmers would benefit from an efficient and validated toolbox to streamline the extraction process and prevent errors in the process. Together, this demonstrates that researchers in cognitive science and computational neuroscience would benefit from a readily-available package for a streamlined extraction of neural network activation.

With `THINGSvision`, we provide a Python toolbox that enables researchers to extract features for most state-of-the-art neural network models for existing or custom image datasets with just a few lines of code. While feature extraction may not seem to be a difficult task for someone with a strong computational background, this toolbox is primarily aimed at supporting those researchers who are inexperienced with Python programming and deep neural network architectures, but interested in the analysis of their representations. However, we believe that even computer scientists will benefit from a publicly available toolbox that is well maintained and efficiently written. Thus, we regard `THINGSvision` as a tool that can be used across research domains.

In the remainder of this article, we introduce and motivate the main functionalities of the library and how to use them. We start by providing an overview of the collection of neural network models for which features can be extracted. The code for `THINGSvision` is publicly available and readily available as a Python package under the MIT license https://github.com/ViCCo-Group/THINGSvision.

### 1.1 Model collection

All neural network models that are part of `THINGSvision` are built in `PyTorch` (Paszke et al., 2019) or `TensorFlow` (Abadi et al., 2015), which are the two most commonly used deep learning frameworks. We include every neural network model that is part of `PyTorch`’s publicly available model-zoo,[torchvision](https://pytorch.org/vision/0.8/models.html), and `TensorFlow`’s model zoo, including many DNN models commonly used in research such as AlexNet (Krizhevsky, Sutskever, & Hinton, 2012), VGG-16 and VGG-19 (Simonyan & Zisserman, 2015), and ResNet (He, Zhang, Ren, & Sun, 2016). Whenever a new vision architecture is added to `torchvision` or `TensorFlow`’s model zoo, `THINGSvision` is designed to automatically make it available, as well.

In addition to models from the `torchvision` and `TensorFlow` library, we provide both feedforward and recurrent variants of CORnet, a recent DNN model that was inspired by the architecture of the non-human primate visual system and that leverages recurrence to more closely resemble biological processing mechanisms (Kubilius et al., 2019, 2018). At the time of writing, CORnet-S is the best performing computational model on the BrainScore benchmark (Schrimpf et al., 2018; Schrimpf, Kubilius, et al., 2020), a composition of various neural and behavioral benchmarks aimed at assessing the degree to which a DNN is a good model of cortical visual object processing.

Moreover, we include both versions of CLIP (Radford et al., 2021), a multimodal DNN model developed by OpenAI that is based on the Transformer architecture (Vaswani et al., 2017), which has surpassed the performance of previous recurrent and convolutional neural networks on a wide range of core natural language processing and image recognition tasks. CLIP’s training procedure makes it possible to simultaneously extract both image and text features for visual concepts and their natural language counterparts. CLIP exists as an advanced, multimodal version of ResNet50 (He et al., 2016) and the so-called Vision-Transformer, `ViT` (Dosovitskiy et al., 2021). We additionally provide the possibility to upload model weights pretrained on custom image datasets beyond ImageNet.

To facilitate the reproducibility of computational analyses across research groups and fields, it is crucial to not only make code pertaining to the proposed analysis pipeline publicly available but additionally offer a general and well documented framework that can easily be adopted by others (Esteban et al., 2018; Peng, 2011; Rush, 2018; Van Lissa et al., 2020). This is why we aspired to follow high software engineering principles such as PEP8 guidelines during development. We regard `THINGSvision` as a toolbox that aims at promoting both the interpretability and comparability of research at the intersection of cognitive science, computational neuroscience, and artificial intelligence. Instead of simply providing an unwieldy collection of existing computational models, we decided to focus on models whose functional composition has been demonstrated to be similar to the primate visual system (Kietzmann et al., 2018; Kriegeskorte, 2015) and models that are widely adopted by the research community.

## 2 Method

`THINGSvision` is a toolbox that was written in the high-level programming language Python and, therefore, requires Python version 3.7 or later to be installed on a user’s machine. The toolbox leverages three of the most widely used packages in the context of machine learning research and numerical analysis, namely PyTorch (Paszke et al., 2019), TensorFlow (Abadi et al., 2015) and NumPy (Harris et al., 2020). Since all relevant NumPy operations were made an integral part of `THINGSvision`, it is not necessary to import NumPy or any other Python package explicitly.

To extract features from a neural network model for a custom set of images, users are first required to select a model and additionally define whether the model’s weights were pretrained on ImageNet (Deng et al., 2009; Russakovsky et al., 2015), randomly initialized or whether the user wants to upload weights that were pretrained on a custom image dataset. If the comparison is aimed at investigating the correspondence between learned representations of a model and brain or behavior, we recommend to use pretrained weights. If the comparison is aimed at investigating how architectural constraints alone can lead to similar representations in models and brain or behavior, then representations from randomly initialized weights carry valuable additional information irrespective of learning (Güçlü & van Gerven, 2015; Schrimpf, Blank, et al., 2020; Storrs, Kietzmann, Walther, Mehrer, & Kriegeskorte, 2020; Yamins et al., 2014). Second, input and output folders as well as the number of samples to be processed in parallel in so-called mini-batches are passed to a function that converts the user’s images into an iterator over mini-batches. This data loader subsequently serves as the input to a function that extracts features for the selected module (e.g., the penultimate layer). The above operations are performed with the following lines of code, which essentially encompass the basic flow of `THINGSvisions`’s extraction pipeline

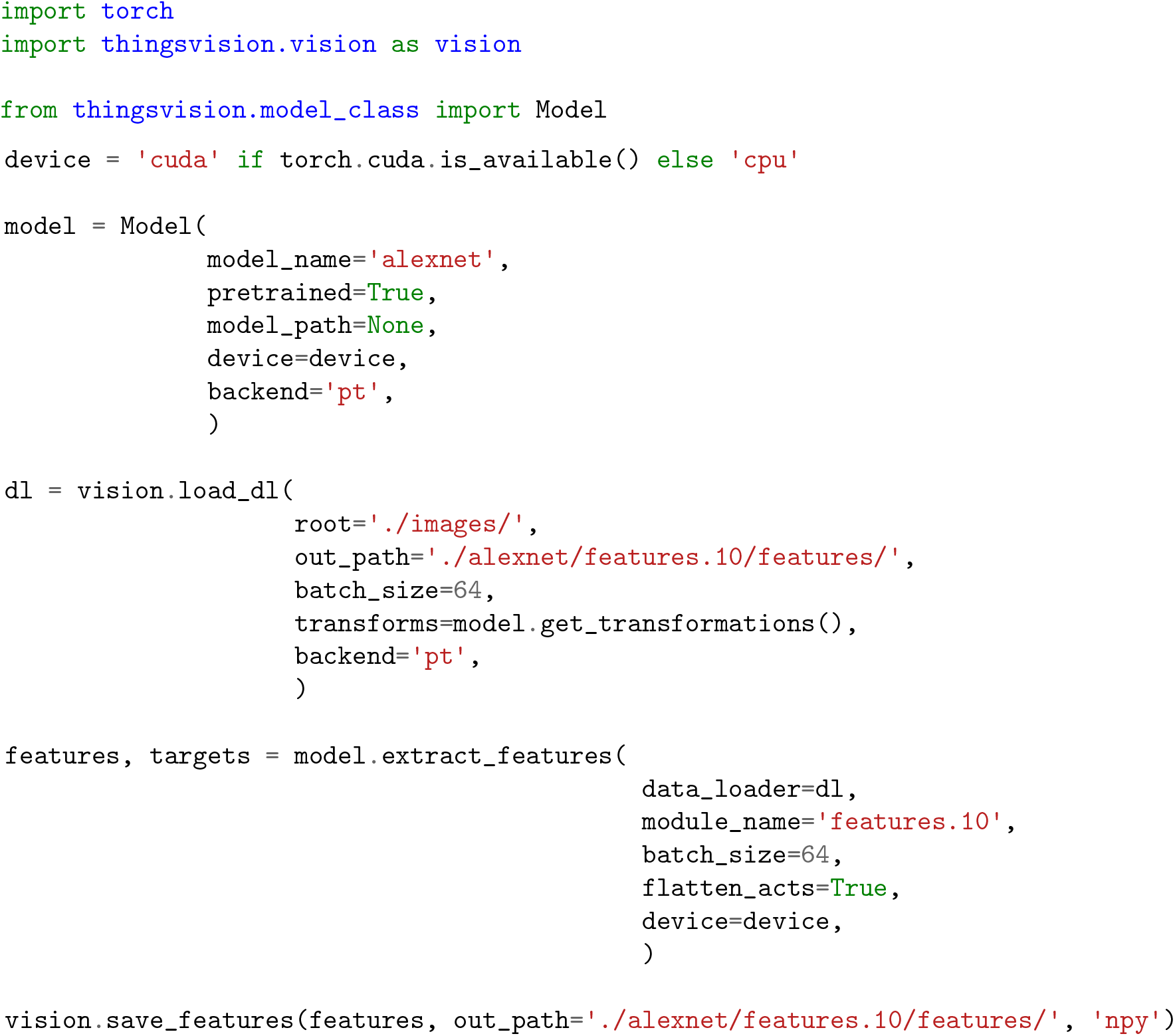

Note that at this point it appears crucial to stress the difference between a layer and a module. Module is a more general reference to the individual parts of a model. A module can refer to non-linearities, pooling operations, batch normalization and convolutional or fully-connected layers, whereas a layer usually refers to an entire model block, such as the composition of the latter set of modules or a single layer (e.g., fully-connected or convolutional). We will, however, use the two terms interchangeably in the remainder of this article whenever a module refers to a layer. Moreover, extracting features is used interchangeably with extracting network activations.

Figure 1 depicts a high-level overview of how feature extraction is streamlined in `THINGSvision`. Given that a user provides the system path to an image dataset, the input to a neural network model is a three-dimensional matrix, 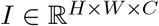, which is the numerical representation of any image. Assuming that a user wants to apply the flattening operation to the activations from the selected module, the output corresponding to each input is a one-dimensional vector, 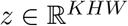.

**Figure 1:**
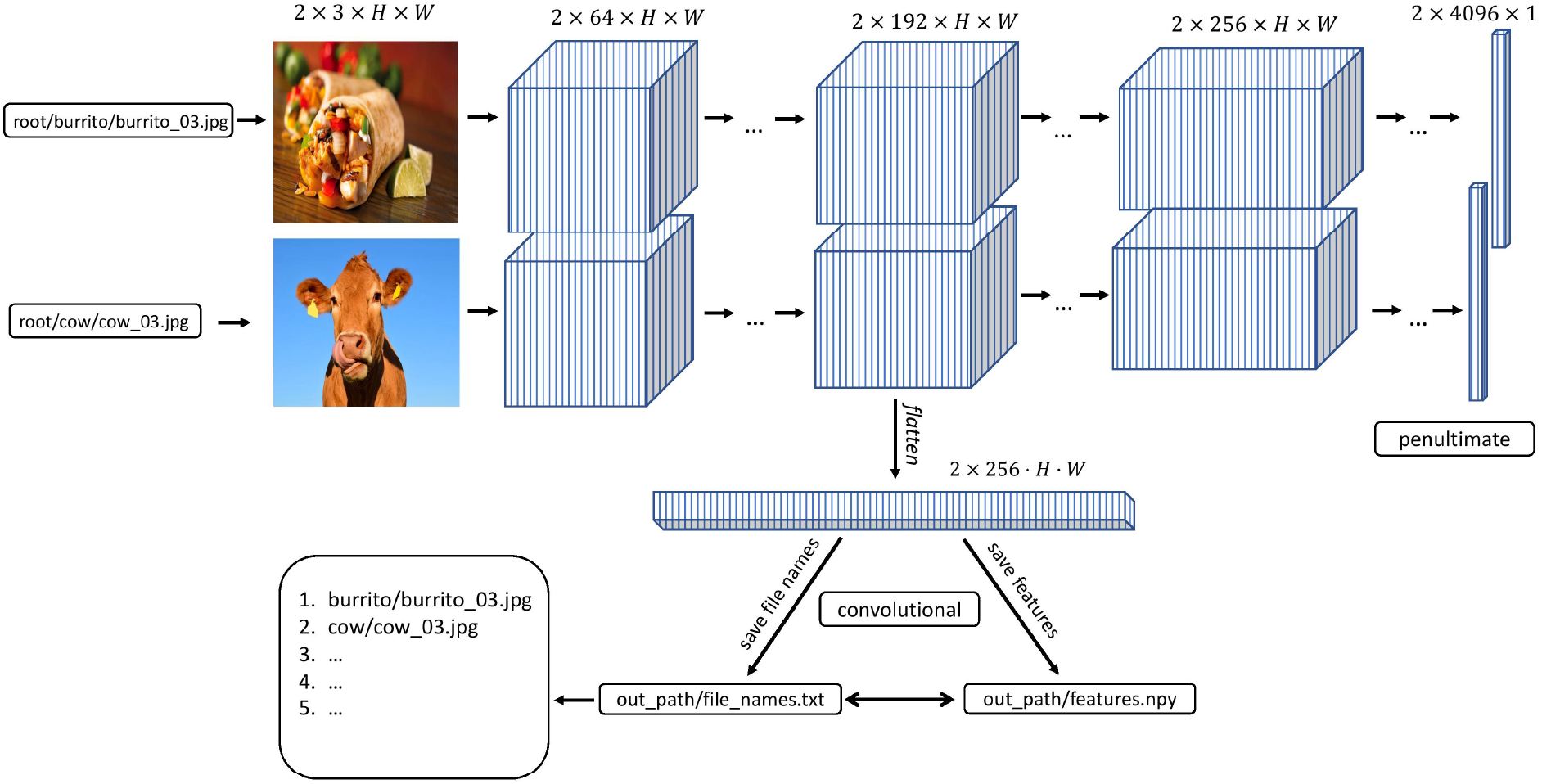
`THINGSvision` feature extraction pipeline for an example convolutional neural network architecture. Images and activations in early layers of the model are represented as four-dimensional arrays. The first dimension represents the batch size, i.e., the number of images in a subsample of the data. For simplicity, in this example this number is set to two. The second dimension refers to the channel-dimension, and the last two dimensions represent the height and width of an image or feature map, respectively.

In the following paragraphs, we will explain both operations and the variables necessary for feature extraction in more detail. We start by introducing variables that we deem helpful for structuring the extraction workflow.

### 2.1 Variables

Before leveraging `THINGSvision`’s full functionality, a user is advised to assign values to seven variables, which, for simplicity, we define as their corresponding keyword argument names: `root`, `model_name`, `pretrained`, `batch_size`, `out_path`, `file_format`, and `device`. Note that this is not a necessity, since the values pertaining to those variables can simply be passed as input arguments to the respective functions. It does, however, facilitate the ease of reading, and in our opinion clearly contributes to a better workflow. Moreover, there is the option to additionally assign a value to the variable `module_name` whose significance we will explain in Section 2.2.2. The above variables, their data types, example assignments, and short descriptions are displayed in Table 1. We will explain the details of these variables in the remainder of this section. We want to stress that our variable assignments are arbitrary examples rather than a general recommendation. The exact values are depending on the specific needs of a user. More advanced users can simply jump to Section 2.2.

**Table 1:**
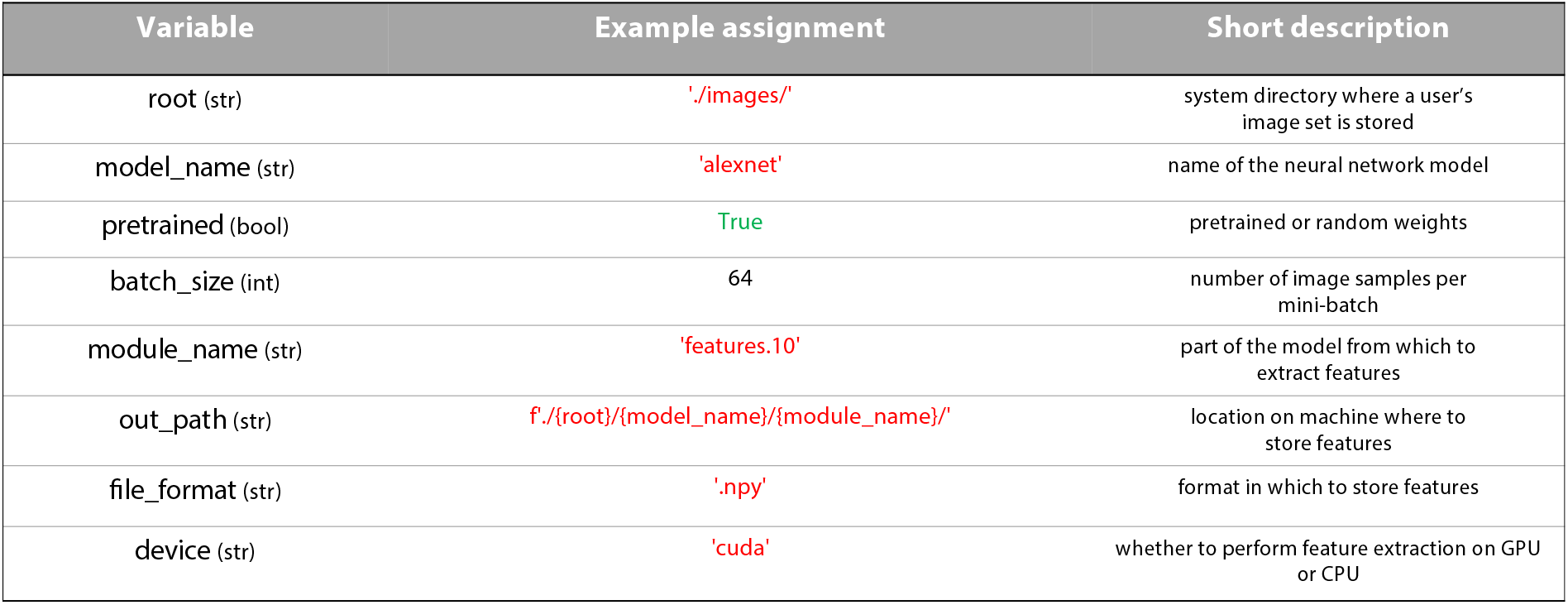
Overview of the variables that are relevant for `THINGSvision`’s feature extraction pipeline and that facilitate a user’s workflow.

#### 2.1.1 Root

We recommend starting with the assignment of the `root` variable. This variable is supposed to correspond to the system directory where a user’s image set is stored.

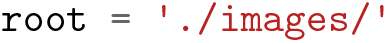

#### 2.1.2 Model name

Next, a user is required to specify the name of the neural network model whose features corresponding to the images in `root` ought to be extracted. The model’s name can be defined as one of the available neural network models in [torchvision](https://pytorch.org/vision/0.8/models.html) or [TensorFlow](https://www.tensorflow.org/api_docs/python/tf/keras/applications). Conveniently, as soon as a new model is added to `torchvision` or `TensorFlow`, it will also be included in `THINGSvision`, since we inherit from both `torchvision` and `TensorFlow`. For simplicity, we use `alexnet` and the `PyTorch` backend throughout the remainder of the article, as shown in Table 1.

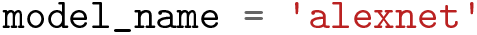

#### 2.1.3 Pretrained

As a subsequent step, a user needs to specify whether to load a pretrained model (i.e., pretrained on `ImageNet`) into memory, or whether to solely load the parameters of a model that has not yet been trained on any publicly available dataset (so-called randomly initialized networks). The latter may be relevant for architectural comparisons when one is concerned not with the knowledge of a model but with its architecture. In the current example, we assume that the user is interested in a model’s knowledge and not its function composition, which is why we set the variable `pretrained` to `true`. Note that `pretrained` must be assigned with a `Boolean` value (see Table 1).

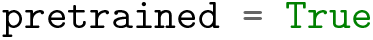

#### 2.1.4 Batch size

Modern neural network architectures process several images at a time in batches. To make the extraction of neural network activations more time efficient, `THINGSvision` follows this processing choice, sampling *B* images in parallel. Thus, the choice of the user lies in the trade-off between processing time and memory usage (GPU memory or RAM). For users who are not concerned with extraction speed, we recommend setting *B* to 32. In our example *B* is set to 64 (see Table 1)

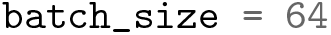

#### 2.1.5 Backend

A user can specify whether to load a neural network model built in `PyTorch` (‘pt’) or `TensorFlow` (‘tf’).

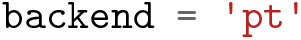

#### 2.1.6 Device

A user can choose between using a CPU and a GPU if a GPU is available. The advantage of leveraging a GPU lies in its faster computation. Note that GPU usage is possible only if a machine is equipped with an NVIDIA GPU.

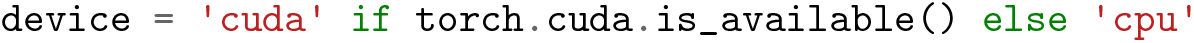

#### 2.1.7 Module name

`Module_name` refers to the part of the model from which network activations should be extracted. In case a user is familiar with the architecture of the neural network model for which features should be extracted, the variable `module_name` can be set manually (e.g., `features.10`). There is, however, the possibility to first inspect the model architecture through an additional function call, and subsequently select a module based on the output of this function. The function prompts a user to select a module, which is then assigned to `module_name` in the form of a string. In Section 2.2.2, we will explain in more detail how this can be done.

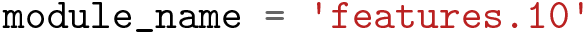

#### 2.1.8 Output directory

Before saving features to disk, a user is required to specify the directory where image features should be stored. For simplicity, in Table 1 we define `out_path` as a succession of previously defined variables.

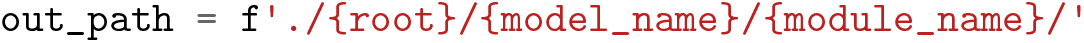

#### 2.1.9 File format

A user can specify the `file_format` in which the image features are stored. This variable can be set either to `hdf5`, `txt`, `mat` or `npy`. If subsequent analyses are performed in Python, we recommend to set `file_format` to `npy`, as storing large matrices in `npy` format is both more memory and time efficient than doing the same in `txt` format. This is due to the fact that the `npy` format was specifically designed to accommodate the storing of large matrices to `NumPy` (Harris et al., 2020).

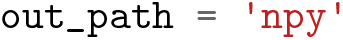

### 2.2 Model & modules

#### 2.2.1 Loading models

With the previously defined variables in place, a user can now start loading a model into a computer’s memory. Since `model_name` is set to `alexnet`, `backend` to `pt` and `pretrained` to `true`, we load an AlexNet model, built in `PyTorch` and pretrained on `ImageNet` into memory with the following line,

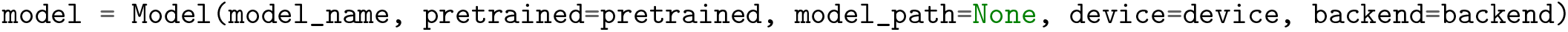

#### 2.2.2 Selecting modules

Before extracting DNN features for an image dataset, a user is required to select the part of the model for which features should be extracted. In case a user is familiar with the architecture of a specific neural network model, they can simply assign a value to the variable `module_name` (see Section 2.1.7). If a user is, however, unfamiliar with the specific architecture of a neural network model, we recommend visualizing the composition of the model’s modules through the following function call,

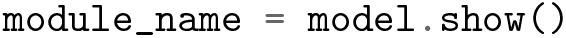

The output of this call, in the case of `alexnet`, looks as follows,

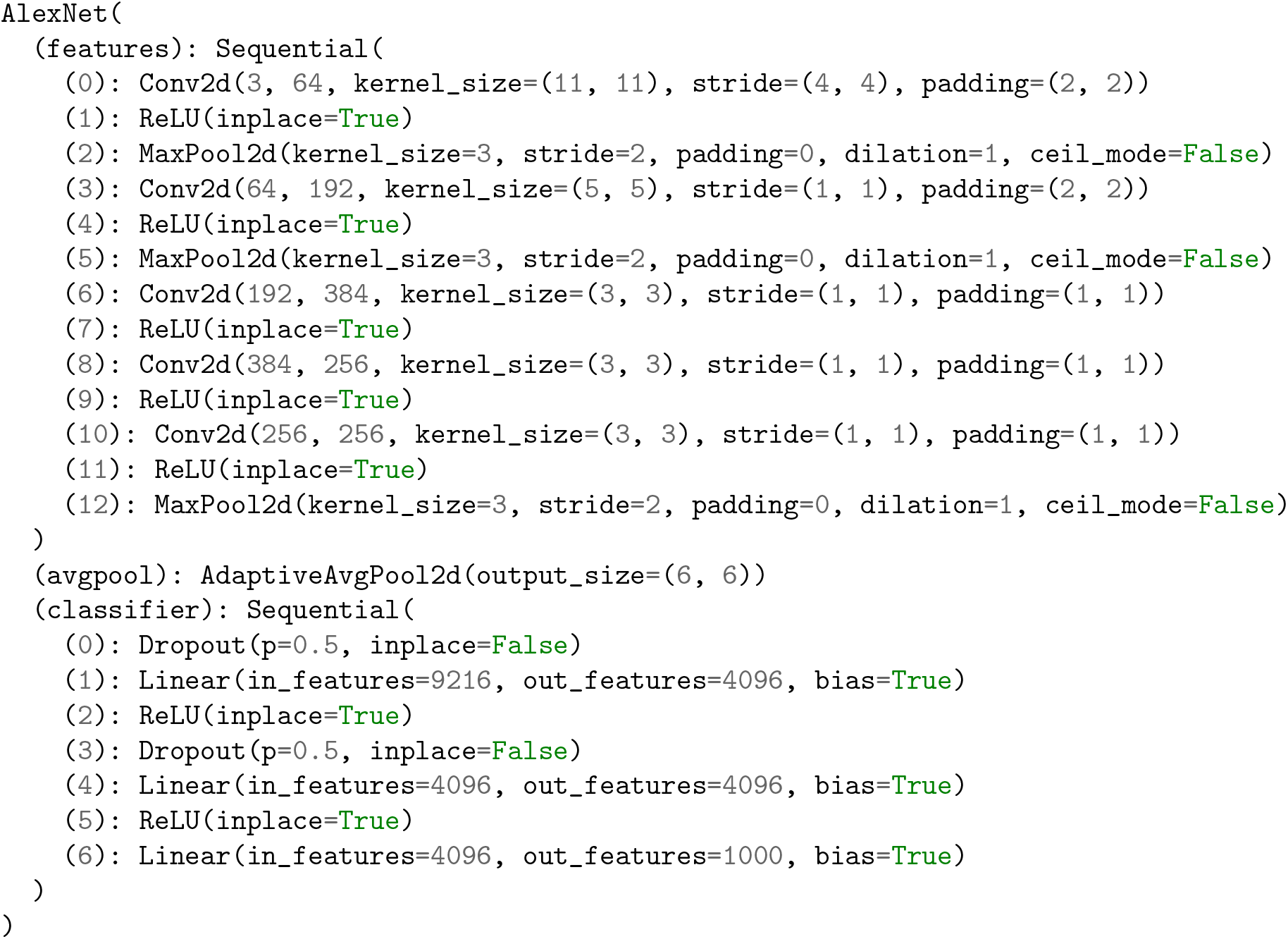

For users unfamiliar with details of neural network architectures, this output may look confusing, given that it is well known that AlexNet consists only of 8 layers. Note, however, that the above terminal output displays the individual modules of AlexNet as well as their specific attributes, such as how many features their inputs and outputs have, or whether a layer is followed by a rectifier non-linearity or pooling operation. Note further that the modules are enumerated in the order in which they appear within the model’s composition. This is crucial for the module selection step. During this step, `THINGSvision` prompts a user to “enter the part of the model for which a user would like to extract image features”. The user’s input is automatically assigned to the variable `module_name` in the form of a string. In order to extract features from layers that correspond to early areas of the primate visual system, we recommend selecting convolutional or pooling modules, and linear layers for later areas that encode high-level features.

It is important to stress that each model in `PyTorch` or `TensorFlow` is represented by a tree structure, where the name of the model refers to the root of the tree (e.g., AlexNet). To access a module, a user is required to compose the string variable `module_name` by both the name of one of the leaves that directly follow the tree’s root (e.g., `features`, `avgpool`, `classifier`) and the number of the module to be selected, separated by a period (e.g., `features.5`). This approach to module selection accounts for all models that are part of `THINGSvision`. How to compose the string variable `module_name` differs between `PyTorch` and `TensorFlow`. We use `PyTorch` module naming.

In this example, we select the 10^th^ module of AlexNet’s leaf `features` (i.e., `features.10`), which corresponds to the fifth convolutional layer in AlexNet (see above). Hence, features will be extracted exclusively for this module.

### 2.3 Dataset & Data loader

Through a dedicated dataset class, `THINGSvision` can deal with various types of image data (`.eps`, `.jpg`, `.jpeg`, `.png`, `.PNG`, `.tif`, `.tiff`) and is able to transform the images into a ready-to-use `PyTorch` or `TensorFlow` dataset. System paths to images can follow the folder structure `./root/class/img_xy.png` or `./root/img_xy.png`, where the former directory contains subfolders for the respective image classes. A dataset is subsequently wrapped with a `PyTorch` or `TensorFlow` iterator to enable batch-wise feature extraction. The above is done with,

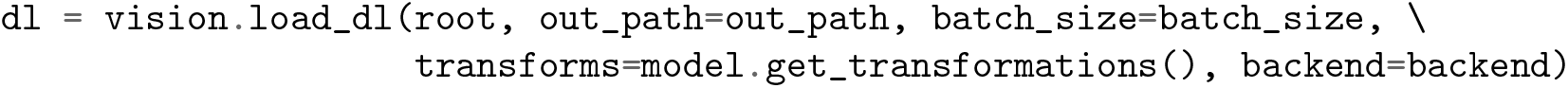

`THINGSvision` automatically sorts image files alphabetically (i.e., A-Z or 0-9). Sorting, however, depends on a machine’s operating system. An alphabetic sort differs across Windows, macOS, and Ubuntu, which is why we provide the possibility to sort the data according to a list of file names, manually defined by the user. The features will, subsequently, be extracted in the order of the provided file names.

This list must follow the `List[str]` data structure (i.e., containing `strings`), such as `[aardvark/aardvark_01.jpg, aardvark/aardvark_02.jpg, …]` or `[aardvark.jpg, anchor.jpg, …]`, depending on whether the dataset tree consists of subfolders for classes (see above). The list of file names can be passed as an optional argument as follows,

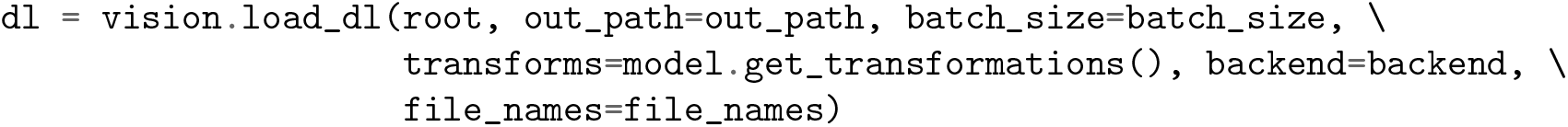

We use the variable `dl` here since it is a commonly used abbreviation for “data loader”. It is, moreover, necessary to pass `out_path` to the above function to save a `txt` to `out_path` consisting of the image names in the order in which features are extracted. This is done to ensure that a user can easily correspond the rows of a feature matrix to the image names, as shown in Figure 1.

### 2.4 Features

The following section is meant for readers curious to understand what is going on under the hood of `THINGSvision`’s feature extraction pipeline and, additionally, who aim to get a better grasp of the dimensions depicted in Figure 1. Readers who are familiar with matrices and tensors may want to skip this section and jump directly to Section 2.4.2, since the following paragraphs are not crucial for using the toolbox. We use mathematical notation to denote images (inputs) and features (outputs).

#### 2.4.1 Extracting features

When all variables necessary for feature extraction are set, the user can extract image features for a specific (here, the fifth convolutional) layer in AlexNet (i.e., `features.10`). Figure 1 shows `THINGSvision`’s feature extraction pipeline for two example images. The algorithm first searches for the images in the `root` folder, subsequently converts them into a ready-to-use dataset, and then passes sub-samples of the data in the form of mini-batches as inputs to the network. For simplicity and to demonstrate the extraction procedure, Figure 1 displays an example of a simplified convolutional neural network architecture. Recall that an image is numerically represented as a three-dimensional array, usually in the following format

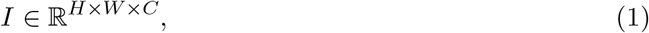

where *H* = height, *W* = width, *C* = channels. *C* = 1 or 3, depending on whether images are represented in grayscale or `RGB` format. In `PyTorch`, however, image batches, denoted as *X*, are represented as four-dimensional tensors,

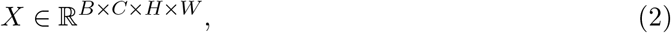

where *B* = `batch_size`, and all other dimensions are permuted. Note, that this is not the case for `TensorFlow`, where image dimensions are not permuted. In the example in Figure 1, *B* = 2, since two images are concurrently processed. The channel dimension, now, represents the tensor’s second dimension (inside the toolbox, it is the first dimension, since Python starts indexing at 0) to more easily apply convolutions to input images. Hence, features at the level of the selected module, denoted as *Z*, are represented as four-dimensional tensors in the format,

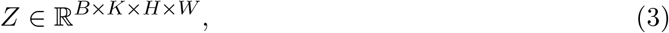

where the channel parameter *C* is replaced with *K* referring to the number of feature maps within a representation. Here, *K* = 256, and *H* and *W* are significantly smaller than at the input level. For most analyses in computational neuroscience, researchers are required to flatten this four-dimensional tensor into a two-dimensional matrix of the format,

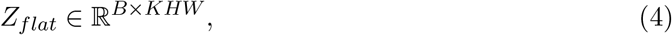

i.e. one vector per image representation in a batch, which is what we demonstrate in the following example. We provide a keyword argument, called `flatten_acts`, that communicates to the function to automatically perform the previous step during feature extraction (see the *flatten* operation in Figure 1). A user must simply set the argument to `True` as follows,

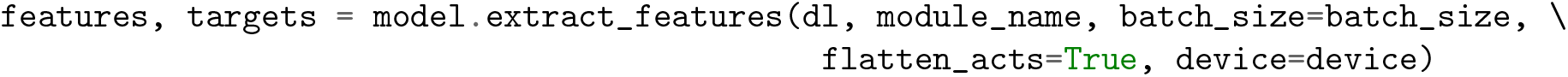

The final, two-dimensional, feature matrix is of the form,

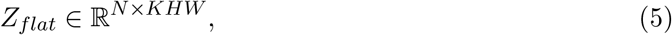

where *N* corresponds to the number of images in the dataset. In addition to the feature matrix, `extract_features` returns a target vector of size *N* × 1 corresponding to the image classes. A user can decide whether to save or ignore this target vector, depending on the subsequent analyses. Note that flattening a tensor is not necessary for feature extraction to work. If a user wants the original four-dimensional tensor, `flatten_acts` must be set to `False`. A flattened representation may be desirable when the neural network representations are supposed to be compared against representations extracted from brain or behavior, which are typically compared using multiple linear regression or by computing correlation coefficients, which cannot operate on multidimensional arrays directly. However, if the goal is to compare activations between different model architectures or leverage interpretability techniques to inspect feature maps, then the tensor should be left in its original four-dimensional shape.

To offer a user more flexibility and control over the feature extraction procedure, we do not provide a default value for this keyword argument. Since a user may want store a four-dimensional tensor in `txt` format to disk, `THINGSvision` comes (1) with a function that slices a four-dimensional tensor into multiple two-dimensional matrices, and (2) a corresponding function that merges the slices back into their original shape at the time of loading the features back into memory.

#### 2.4.2 Saving features

To save network activations (no matter from which part of the model) in a flattened format, the following function can be called,

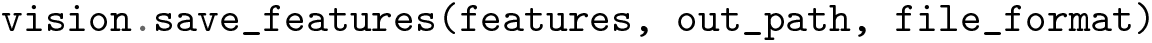

When features are extracted from any of the convolutional layers of the model, the output is a four-dimensional tensor. Since it is not trivial to save four-dimensional tensors in `txt` format to be readily used for subsequent analyses of a model’s feature maps, a user is required to set the file format argument to `hdf5`, `npy`, or `mat`, of which all enable the saving of four-dimensional tensors in their original shape.

When storing network activations from convolutional layers in their flattened format, it is possible to run into `MemoryErrors`. We account for that potential caveat with splitting two-dimensional matrices into *k* equally large splits, whenever that happens. The default value of *k* is set to 10. If 10 splits are not sufficient to counteract the memory issues, a user can change this value to a larger number. We recommend trying multiples of 10, such as

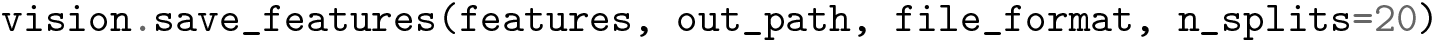

To merge the array splits back into a single, two-dimensional, feature matrix, a user can call,

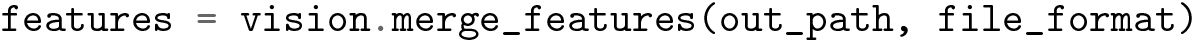

### 2.5 Representational Similarity Analysis

Representational Similarity Analysis (RSA), a technique that originated in cognitive computational neuroscience, can be used to relate object representations from different measurement modalities (e.g., fMRI or behavior) and different computational models with each other (Kriegeskorte, Mur, & Bandettini, 2008; Kriegeskorte, Mur, Ruff, et al., 2008). RSA is based on representational dissimilarity matrices (RDMs), which capture the representational geometry present in a given system (e.g., in the brain or a DNN), thereby abstracting away from the underlying multivariate pattern. Rather than directly comparing measurements, RDMs compare representational similarities between two systems. RDMs are symmetric, square matrices, where the rows and columns are indexed by the different conditions or objects. Hence, RSA is a convenient analysis tool to compare visual object representations obtained from different DNNs.

The dissimilarity between each object pair (e.g., two images) is computed within the row space of an RDM. Dissimilarity is quantified as the distance between two objects in the measured representational space, defined by the chosen distance metric. The user can choose between the Euclidean distance (`euclidean`), the correlation distance (`correlation`), the cosine distance (`cosine`) and a radial basis function applied to pairwise distances (`gaussian`). Equivalent object representations show a dissimilarity score close to 0. For the `correlation` and `cosine` distances, the maximum dissimilarity score is bounded to 2, whereas there is no theoretical upper limit for the `euclidean` distance.

Since RDMs are symmetric around their main diagonal, it is simple to compare them by correlating their lower or upper triangles. We include both the possibility to compute and visualize an RDM and to correlate the upper triangles of two distinct RDMs. Computing an RDM based on a Pearson correlation distance matrix is as simple as calling

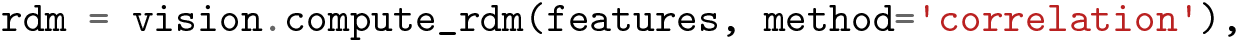

Note that similarities are computed between conditions or objects, not features. To compute the representational similarity between two distinct RDMs, a user can make the following call,

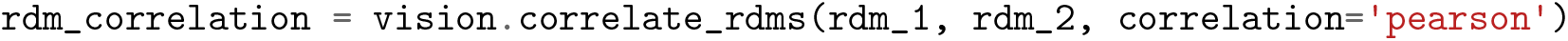

The default `correlation` value is the Pearson correlation coefficient, but this can be changed to `spearman` if a user assumes that the similarities are not ratio scale and require the computation of a Spearman rank correlation (Arbuckle, Yokoi, Pruszynski, & Diedrichsen, 2019; Nili et al., 2014). To visualize an RDM and automatically save the output image (in `.png` or `.jpg` format) to disk, one may call

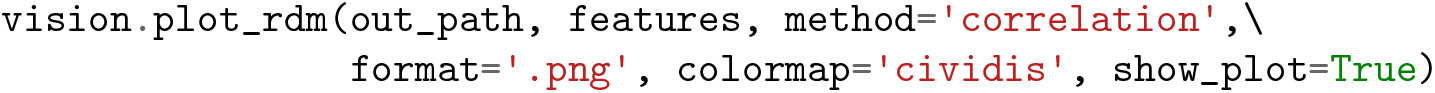

The default value of `format` is set to `.png` but can easily be changed to `.jpg`. Note that `.jpg` is a lossy image compression, whereas `.png` is lossless, and, hence, with `.png` no information gets lost during compression. Therefore, the `format` argument influences both the size and the final resolution of the RDM image representation. The `dpi` value is set to 200 to guarantee for a high image resolution, even if `.jpg` is selected.

## 3 Results & Applications

To demonstrate the usefulness of `THINGSvision`, in the following, we present analyses of the image representations of different deep neural network architectures and compare them against representations obtained from behavioral experiments (3.1.1) and functional MRI responses to higher visual cortex (3.1.2). To qualitatively inspect the DNN representations, we compute and visualize representational dissimilarity matrices (RDMs) within the framework of representational similarity analysis (RSA), as introduced in Section 2.5. Moreover, we calculate the Pearson correlation coefficients between human and DNN representations to quantify their similarities, and show how this can easily be done with `THINGSvision`. We measure the correspondence between layer activations and human brain or behavioral representations as the Pearson’s correlation coefficient, in line with the recent finding that the linearity assumption holds for functional MRI data which validates the use of an interval rather than an ordinal scale (Arbuckle et al., 2019).

In addition to results for pretrained models, we compare randomly initialized models against human brain and behavioral representations. This reveals the degree to which the architecture by itself, without any prior knowledge (e.g. through training), may perform above chance and which model achieves the highest correspondence to behavioral or brain representations under these circumstances. Indeed, a comparison to randomly-initialized networks is increasingly used as a baseline for comparisons(e.g., Cichy, Khosla, Pantazis, Torralba, & Oliva, 2016; Güçlü & van Gerven, 2015; Schrimpf, Blank, et al., 2020; Storrs, Kietzmann, et al., 2020; Yamins et al., 2014).

Note that this section should not be regarded as an investigation in its own right. It is supposed to demonstrate the usefulness and versatility of the toolbox. This is the main reason for why we do not make any claims about hypotheses and how to test them. RSA is just one out of many potential applications, of which a subset is mentioned in the Discussion section.

### 3.1 The penultimate layer

The correspondence of a DNN’s penultimate layer to human behavioral representations has been studied extensively and is therefore often used when investigating the representations of abstract visual concepts in neural network models (e.g., Bankson et al., 2018; Battleday, Peterson, & Griffiths, 2019; Cichy, Kriegeskorte, Jozwik, van den Bosch, & Charest, 2019; Jozwik, Kriegeskorte, Cichy, & Mur, 2018; Mur et al., 2013; Peterson, Abbott, & Griffiths, 2018). To the best of our knowledge, our study is the first to compare visual object representations extracted from CLIP (Radford et al., 2021) against the representations of well known vision models that have previously shown a close correspondence to neural recordings of the primate visual system. We computed RDMs based on the Pearson correlation distance for seven models, namely AlexNet (Krizhevsky et al., 2012), VGG16 and VGG19 with batch normalization (Simonyan & Zisserman, 2015), which show a close correspondence to brain and behavior (Schrimpf et al., 2018; Schrimpf, Kubilius, et al., 2020), ResNet50 (He et al., 2016), BrainScore’s current leader CORnet-S (Kubilius et al., 2019, 2018; Schrimpf, Kubilius, et al., 2020), and OpenAI’s CLIP variants CLIP-RN and CLIP-ViT (Radford et al., 2021). The comparison was done for six different image datasets that included functional MRI of the human visual system and behavior (Bankson et al., 2018; Cichy et al., 2019; Hebart et al., 2020; Mohsenzadeh et al., 2019; Mur et al., 2013). For the neuroimaging datasets, participants viewed different images of objects while performing an oddball detection task in an MRI scanner. For the behavioral datasets, participants completed similarity judgments using the multiarrangement task Bankson et al. (2018); Mur et al. (2013) or a triplet odd-one-out task Hebart et al. (2020).

Note that Bankson et al. (2018) exploited two different datasets which we label with “(1)” and “(2)” in Figure 2. The number of images per dataset are as follows: (Cichy, Pantazis, & Oliva, 2014; Kriegeskorte, Mur, Ruff, et al., 2008; Mur et al., 2013): 92; (Bankson et al., 2018) 84 each; (Cichy et al., 2016, 2019): 118; (Mohsenzadeh et al., 2019): 156; (Hebart et al., 2019, 2020): 1854. For each of these datasets except for Mohsenzadeh et al. (2019), we additionally computed RDMs for group averages obtained from behavioral experiments. Furthermore, we computed RDMs for brain voxel activities obtained from fMRI recordings for the datasets used in Cichy et al. (2014), Cichy et al. (2016), and Mohsenzadeh et al. (2019), based on voxels inside a mask covering higher visual cortex.

**Figure 2:**
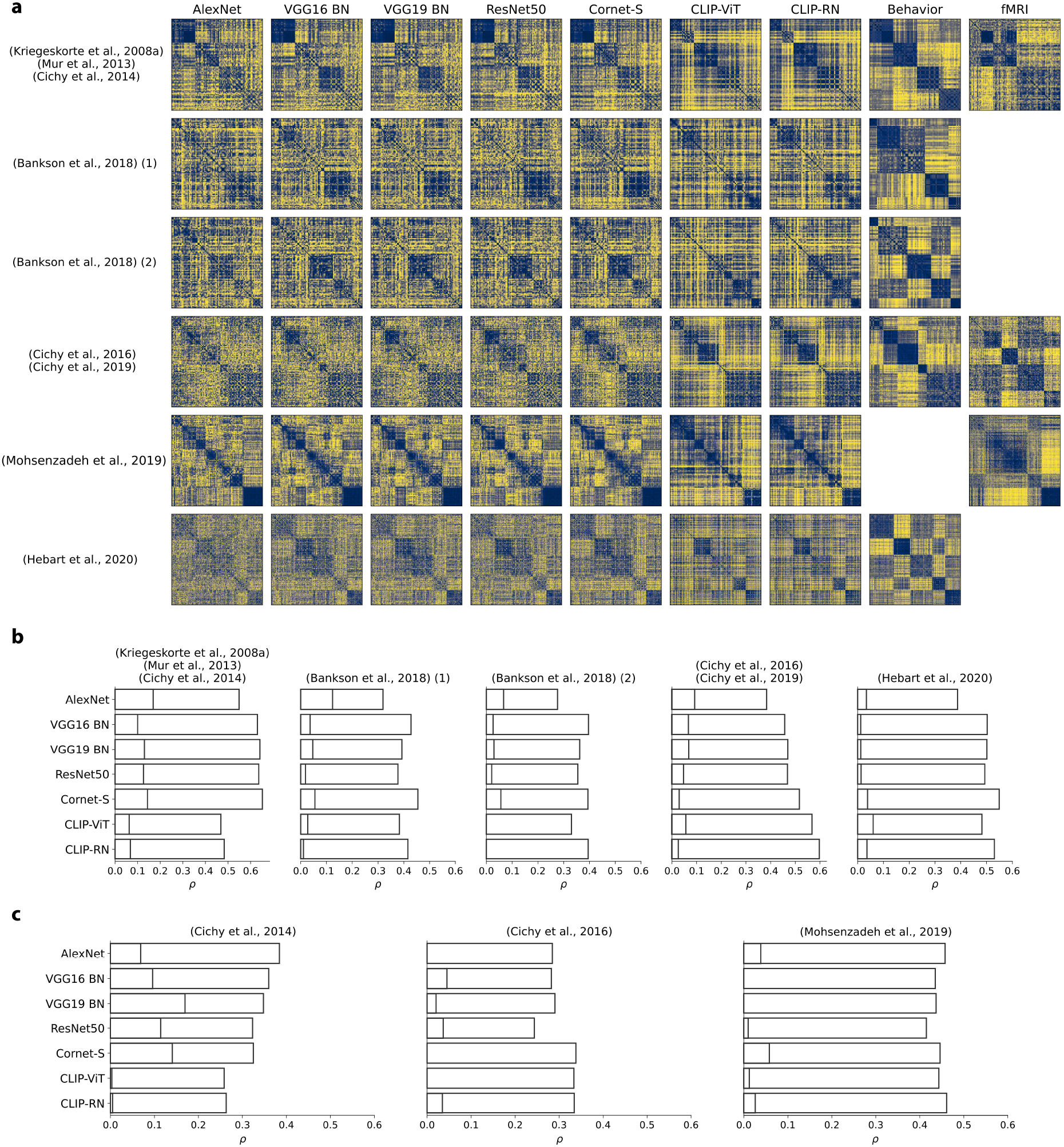
**a**. RDMs for penultimate layer representations of different pretrained neural network models, for group averages of behavioral judgments, and for fMRI responses to higher visual cortex. For Mohsenzadeh et al. (2019), no behavioral experiments had been conducted. For both datasets in Bankson et al. (2018), and for Hebart et al. (2020), no fMRI recordings were available. For display purposes, Hebart et al. (2020) was downsampled to 200 conditions. RDMs were reordered according to an unsupervised clustering. **b/c**. Pearson correlation coefficients for comparisons between neural network representations extracted from the penultimate layer and behavioral representations (b) and representations corresponding to fMRI responses of higher visual cortex (c). Activations were extracted from pretrained and randomly initialized models.

Figure 2 (a) visualizes all RDMs. We clustered RDMs pertaining to group averages of behavioral judgments into five object clusters and sorted the RDMs corresponding to object representations extracted from DNNs according to the obtained cluster labels. The image datasets used in Cichy et al. (2014); Kriegeskorte, Mur, Ruff, et al. (2008); Mur et al. (2013) and Mohsenzadeh et al. (2019) were already sorted according to object categories, which is why we did not perform a clustering on RDMs for those datasets. The number of clusters was chosen arbitrarily. The reordering was done to highlight the similarities and differences in RDMs.

#### 3.1.1 Behavioral correspondences

##### Pretrained weights

Across all compared DNN models, CORnet-S and CLIP-RN showed the overall closest correspondence to behavioral representations. CORnet-S, however, was the only model that performed well across all datasets. CLIP-RN showed a high Pearson correlation (ranging from 0.40 to 0.60) with behavioral representations across most datasets, with Mur et al. (2013) being the only exception, for which both CLIP versions performed poorly. Interestingly, for one of the datasets in Bankson et al. (2018), VGG16 with batch normalization (Simonyan & Zisserman, 2015) outperformed both CORnet-S and CLIP-RN (see Figure 2 (b)). AlexNet consistently performed the worst for behavioral fits. Note that the broadest coverage of visual stimuli is provided by Hebart et al. (2019, 2020), which should therefore be seen as the most representative result (rightmost column in Figure 2 (b)).

##### Random weights

Another interesting finding is that for randomly-initialized weights, CLIP-RN is the poorest performing model in four out of five datasets (see bars in Figure 2 (b) corresponding to lower correlation coefficients). Here, AlexNet seems to be the best performing model across datasets, although it achieved the lowest correspondence to behavioral representations when leveraging a pretrained version (see Figure 2 (b))). This indicates the possibility of complex interactions between model architectures and training objectives that require further investigations which `THINGSvision` may facilitate.

#### 3.1.2 Brain correspondences

We performed a similar analysis as above, but this time leveraging RDMs corresponding to fMRI responses to examine the correlation between model and brain representations of higher visual cortex. We first report results obtained from analyses with pretrained models.

##### Pretrained weights

While AlexNet (Krizhevsky et al., 2012) showed the worst correspondence to human behavior in four out of five datasets (see Figure 2 (c)), AlexNet correlated strongly with representations extracted from fMRI responses to higher visual cortex, except for the dataset used in Cichy et al. (2016) (see Figure 2 (c)). This is interesting, given that among the entire set of analyzed deep neural network models AlexNet shows the poorest performance on ImageNet (Russakovsky et al., 2015). This result contradicts findings from previous studies arguing that object recognition performance is correlated with correspondences to fMRI recordings (Schrimpf, Kubilius, et al., 2020; Yamins et al., 2014). This time, CORnet-S and CLIP-RN performed well for the datasets used in Cichy et al. (2016) and in Mohsenzadeh et al. (2019), but were among the poorest performing DNNs for Cichy et al. (2014). Note, however, that the dataset used in Cichy et al. (2014) is highly structured and contains a large number of faces and similar images, something AlexNet might pick up more easily in its image features but something that is not reflected in human behavior (Grootswagers & Robinson, 2021).

##### Random weights

When comparing representations corresponding to network activations from models with random weights, there appears to be no consistent pattern as to which model correlated most strongly with brain representations of higher visual cortex, although VGG16 and CORnet-S were the only two models that yielded a Pearson correlation coefficient > 0 across datasets. Note, however, that for each model we extracted network activations from the penultimate layer. Results might look different when extracting activations from earlier layers of the networks or when reweighting the DNN features prior to RSA (Kaniuth & Hebart, 2020; Storrs, Khaligh-Razavi, & Kriegeskorte, 2020). We leave further investigations to future studies, as our analyses should only demonstrate the applicability of our toolbox.

#### 3.1.3 Model comparison

Although CORnet-S and CLIP-RN achieved the overall highest correspondence to both behavioral and human brain representations, our results indicate that more recent, deeper neural network models are not necessarily preferred over previous, shallower models, at least when exclusively leveraging the penultimate layer of a network. Their correspondences appear to be highly dataset-dependent. Although a pretrained version of AlexNet correlated poorly with representations obtained from behavioral experiments (see Figure 2 (b)), there are datasets where AlexNet showed close correspondence to brain representations (see Figure 2 (c)). Similarly, VGG16 was mostly outperformed by CLIP-RN, but in one out of five datasets it yielded a higher correlation with behavioral representations than CLIP-RN.

## 4 Discussion

Here we introduce `THINGSvision`, a Python toolbox for extracting activations from hidden layers of a wide range of deep neural network models. We designed `THINGSvision` to facilitate research at the intersection of cognitive science, computational neuroscience, and artificial intelligence.

Recently, an API was released (Mehrer, Spoerer, Jones, Kriegeskorte, & Kietzmann, 2021) that enables the extraction of image features from AlexNet and vNet without the requirement to install any library, making it a highly user-friendly contribution to the field. Apart from requiring an installation of Python, `THINGSvision` provides a comparably simple way to extract network activations, yet for a much broader set of DNNs, for `PyTorch` and `TensorFlow` backends, and with a higher degree of flexibility and control over the extraction procedure. `THINGSvision` can easily be integrated with any other computational analysis pipeline performed in Python or Matlab. We additionally allow for a streamlined comparison of visual object representations obtained from various DNNs employing representational similarity analysis.

We demonstrated the usefulness of `THINGSvision` through the application of RSA and the quantification of correspondences between representations extracted from models and human behavior (or brains). Please note that the extracted network activations are not only useful for visualizing and comparing network activations through frameworks such as RSA, but for any downstream application, including regression onto brain data, (Güçlü & van Gerven, 2015; Yamins et al., 2014), feature selectivity analysis (e.g., Xu, Zhang, Zhen, & Liu, 2021), or fine-tuning of individual layers for external tasks (e.g., Khaligh-Razavi & Kriegeskorte, 2014; Tajbakhsh et al., 2016).

`THINGSvision` enabled us to investigate object representations of CLIP (Radford et al., 2021) against representations extracted from other neural network models as well as representations from behavioral experiments and fMRI responses to higher visual cortex. To understand why Transformer layers and multimodal training objectives help to achieve strong correspondences to behavioral representations (see Figure 2 (b)), further studies are encouraged to investigate the representations of CLIP and its differences to previous DNN architectures with unimodal objectives.

We hope that `THINGSvision` will serve as a useful tool that supports researchers in carrying out such analyses, and we intend to extend the set of models and functionalities that are integral to `THINGSvision` over the coming years as a function of advancements and demands in the field.

## Author Contributions

LM designed the toolbox. LM programmed the software. LM and MNH collected the data. LM and MNH analyzed and visualized the data. MNH supervised the study. MNH acquired funding. LM and MNH wrote the manuscript. All authors agreed with the final version of the manuscript.

## Acknowledgments

The authors would like to thank Katja Seeliger, Oliver Contier and Philipp Kaniuth for useful comments on earlier versions of this paper, and in particular Hannes Hansen, who helped running all sorts of tests and enhancing continuous integration of the toolbox.

## A Appendix

In the code example below, we demonstrate both the flexibility and ease of use of our toolbox compared to using solely PyTorch. THINGSvision is more versatile and does not require the user to be adept in PyTorch, whereas when using PyTorch more knowledge about neural network architectures and tensor dimensions is crucial. This might not cause difficulties for someone experienced in Python programming and Machine Learning, but is not trivial for researchers who are not as familiar with this area of Computer Science.

### THINGSvision

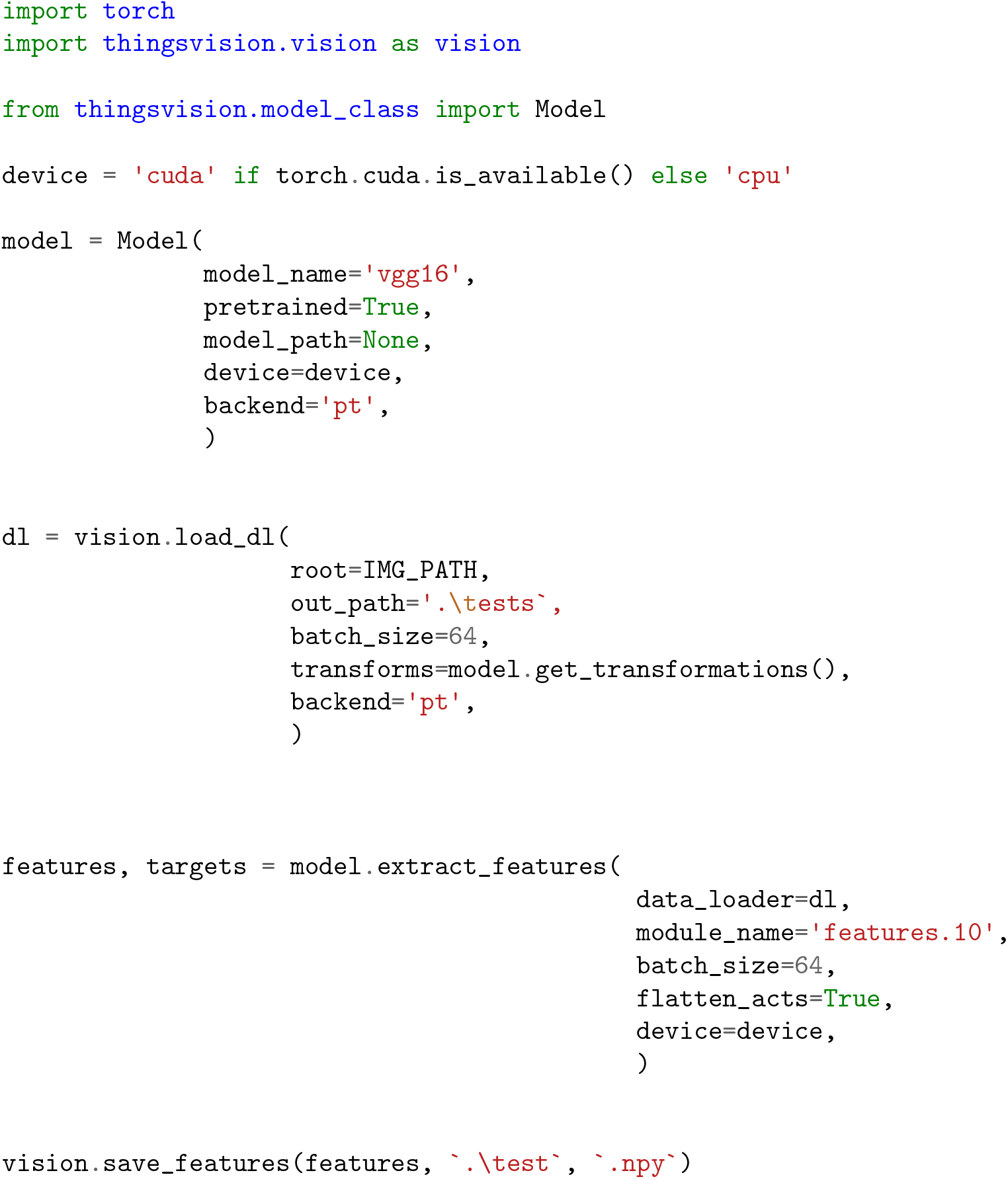

### PyTorch

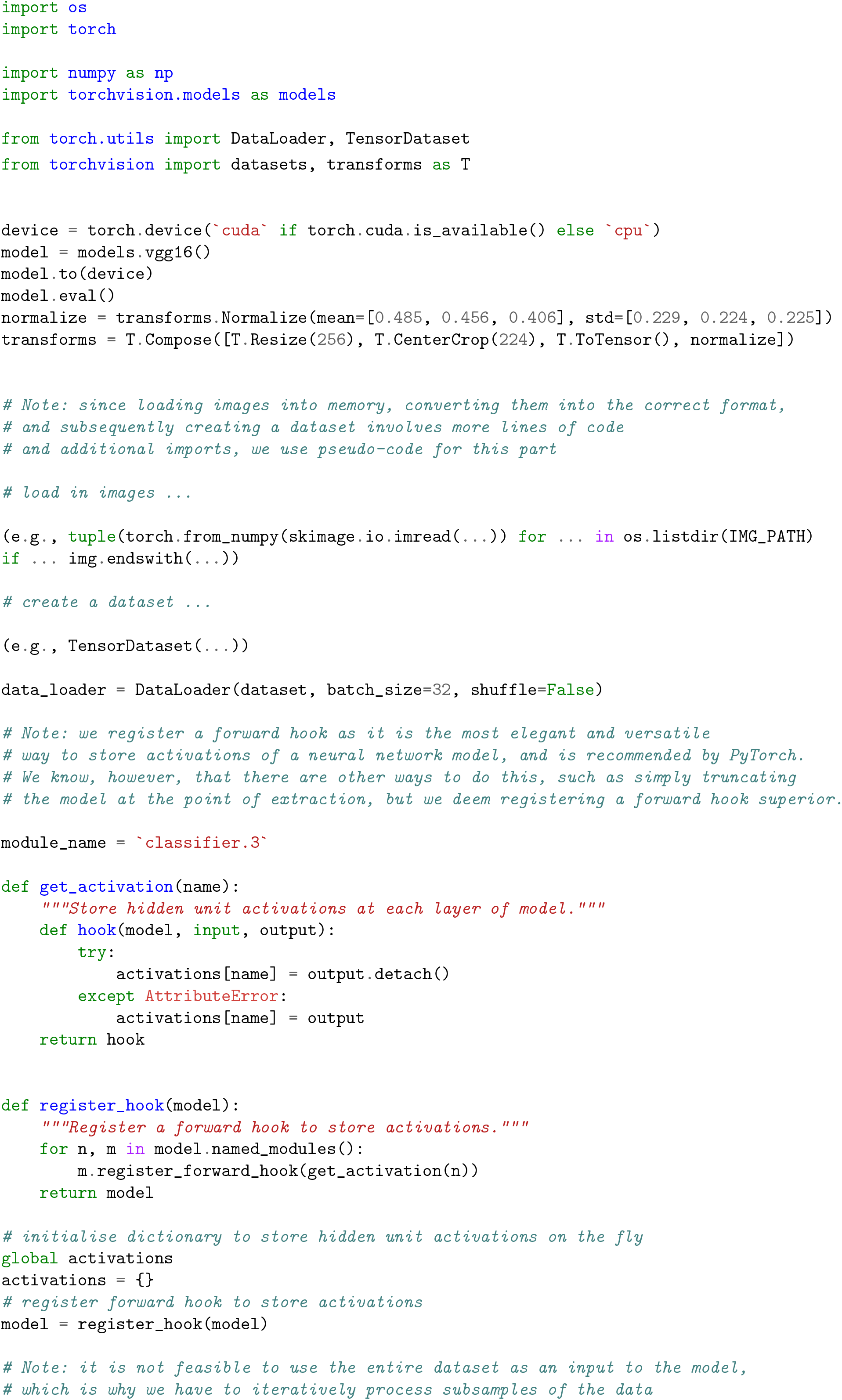
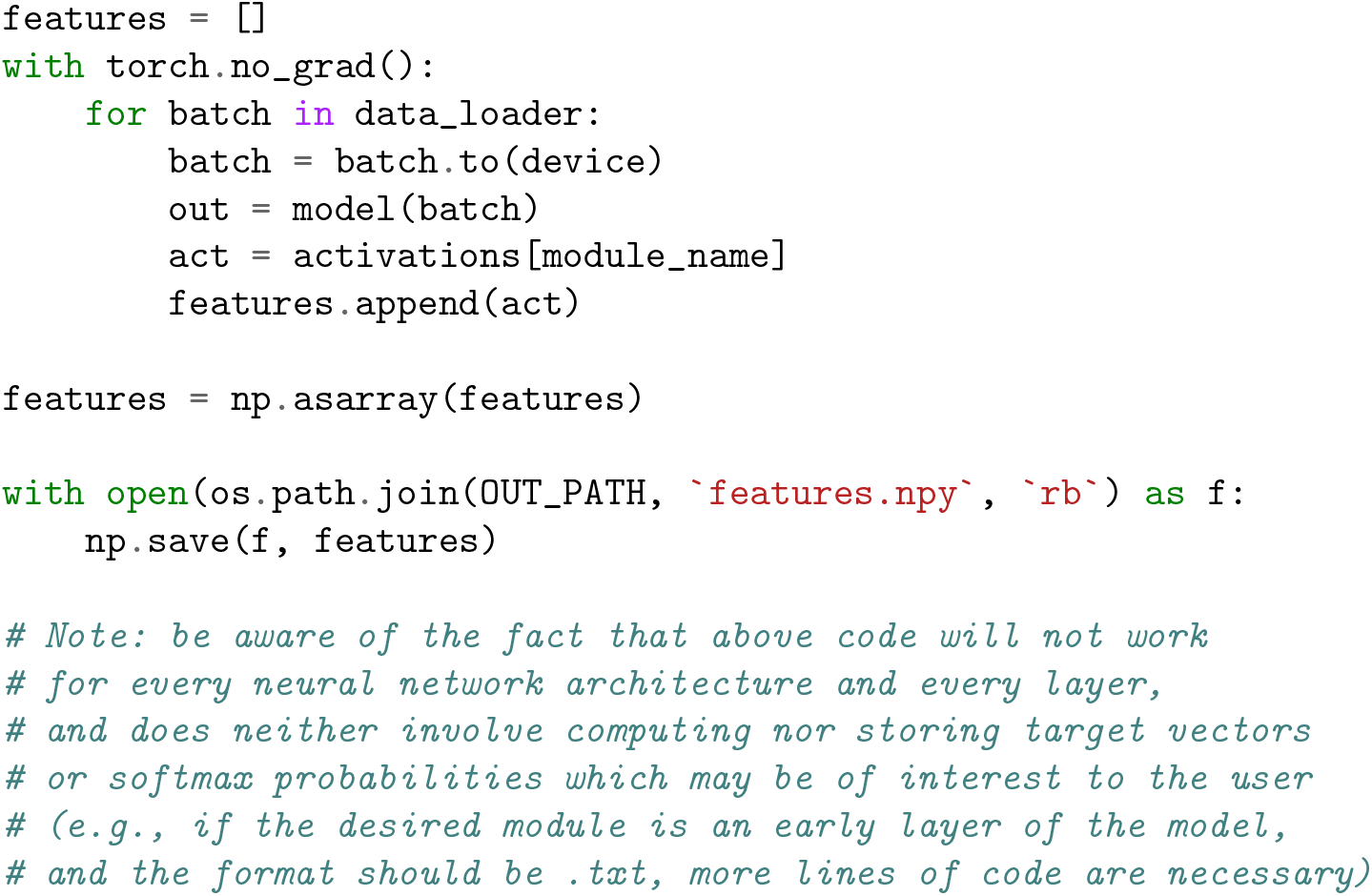

